# Calcium-dependent cytoskeletal collapse and recovery of axons after partial laser ablation

**DOI:** 10.1101/2025.08.08.668418

**Authors:** Ashish Mishra, Pooja Joshi, Md Arsalan Ashraf, Pramod Pullarkat

## Abstract

Traumatic stretch or crush injury to axons causes widespread and often irreversible damage to the axonal cytoskeleton, in which calcium-mediated breakdown is known to play a central role. Unlike complete transection, where recovery must proceed through formation of a new growth cone, milder injury can disrupt the axonal cytoskeleton while leaving the plasma membrane intact. How the cytoskeleton fails, and how it can recover, under these conditions remains unclear. Here we address this using a partial laser-ablation method that damages the cytoskeleton and evokes a calcium transient while preserving membrane continuity. We show that the ensuing cytoskeletal retraction is set by a mechanical balance between acto-myosin contractility and microtubule stability: stabilizing microtubules or inhibiting acto-myosin contractility suppresses retraction. Moreover, chelating extracellular calcium mitigates degeneration and, in a subset of axons, permits complete recovery. We also show that microtubules and actin filaments show distinct loss and recovery dynamics and provide a hypothesis for the “burning-fuse” –like depolymerization of the microtubule bundle. These findings provide insights into how the axonal cytoskeleton collapses and recovers after injury and suggest strategies for mitigating damage.

## Introduction

Neuronal cells are highly susceptible to damage caused by mechanical stress (Hill et al., 2016). Impacts or sudden acceleration of the head can lead to concussion, diffuse axonal injury (DAI), or traumatic brain injury (TBI) (Hill et al., 2016; Johnson et al., 2013; Smith et al., 2003). Axons of the brain are particularly vulnerable as shear/stretch deformation of the brain leads to stretching of the axons that form intricate networks within the brain (Keating & Cullen, 2021; Tang-Schomer et al., 2010). Injuries to the spinal cord or to the peripheral nerves are also common, either due to local compression or local stretch during accidents, excessive limb movement, dislocation, breech delivery, or during surgical manipulation (Anthes et al., 1995; Aoki, 2005; Avis & Power, 2018; Fournier et al., 2015; Sharp et al., 2021; Vialle et al., 2007; Yap et al., 2017). Such injuries form a leading cause of death and debilitating disabilities, especially among the young (Karaboue et al., 2024).

Stretch or compression-induced damage to axons can occur either due to the direct disruption of axonal cytoskeletal elements or structural degradation triggered by molecular events (Büki & Povlishock, 2006; Fournier et al., 2015; Tang-Schomer et al., 2010). Calcium influx into the axons is a key event that is well known to result in widespread axonal damage after injury (Büki & Povlishock, 2006). This influx can occur via the opening of stretch-sensitive ion channels, reversal of Na^+^-Ca^++^ exchanger, voltage-gated ion channels, or via pore formation in the axonal membrane (Aydın et al., 2023; Khaitin, 2021; Wolf et al., 2001). Calcium entry can trigger a series of cascade events which drastically augment damage (Büki & Povlishock, 2006). It triggers the release of calcium from major internal stores like the endoplasmic reticulum, and this calcium induced calcium release spreads rapidly along the axon away from the site of injury (Villegas et al., 2014; Ziv & Spira, 1995). Following this short timescale molecular event, elevated calcium activates proteases like calpain. Calpain cleaves cytoskeletal proteins such as spectrin, neurofilaments, and microtubule-associated proteins, to initiate a slower and more widespread phase of cytoskeletal degradation within the axon (Büki & Povlishock, 2006; Heller et al., 2025b; Ma, 2013; Vosler et al., 2008). Other mechanisms like mitochondrial failure and oxidative stress can amplify damage (Frati et al., 2017; Pivovarova & Andrews, 2010). This degradation disrupts axonal transport, causes axonal swelling, and ultimately leads to complete degeneration of the axon (Frati et al., 2017).

Understanding cytoskeletal stability post-injury, and calcium’s role in this offers insights into potential therapeutic strategies to limit secondary injury. Currently, treatment strategies targeting calcium dysregulation, calpain activation, and oxidative stress have emerged as promising approaches to mitigate neuronal damage in conditions like diffuse axonal injury and nerve stretch injury. However, a complete understanding of the molecular mechanisms and cytoskeletal alterations that occur post-injury and potential recovery mechanisms post-treatment is still lacking. Current laboratory methods include using of stretchable substrates (Tang-Schomer et al., 2010), microfluidics (Pan et al., 2022), and manipulation or transection of axons using mechanical devices (Blizzard et al., 2007; Gallo, 2004; Kerschensteiner et al., 2005; Shao et al., 2019) or using lasers (Cengiz et al., 2012; Kunik et al., 2011). Although such techniques have greatly improved our understanding of axonal degeneration, several aspects of axonal stability after injury and especially the potential recovery pathways remain poorly explained.

Here, we present a partial laser ablation method which can induce local damage to the cytoskeleton of cultured chick Dorsal Root Ganglia axons and trigger a calcium response while maintaining the axonal plasma membrane connectivity. We show that such damage results in a catastrophic loss of cytoskeletal support to the axon, resulting in complete axonal atrophy. However, axonal degradation can be mitigated either by stabilizing microtubules or by destabilizing acto-myosin complex. Conversely, microtubule destabilization or actin stabilization hastens retraction. Almost complete inhibition is achieved in combination treatment. Equally remarkably, a subset of axons exhibits complete recovery when extracellular calcium is chelated, with actin filaments and microtubules showing starkly different recovery patterns. We contrast this recovery pattern with that seen in complete transection, where a growth cone needs to be regenerated. Further, we show that recovery of actin filaments alone does not guarantee axonal recovery, whereas microtubule regrowth can lead to full recovery of damaged axons. Finally, we hypothesize on a possible mechanism to explain the degeneration pattern observed for the axonal microtubule bundle.

## Materials and Methods

### Cell culture medium

L-15 medium (21083–027, Thermo Fisher Scientific) was made viscous using autoclaved methylcellulose (H7509-100 g; Sigma-Aldrich, Darmstadt, Germany) at a ratio of 100 ml to 0.6 g by stirring overnight at 4 ^∘^C. This was then supplemented with 10% (v/v) heat-inactivated fetal bovine serum (10100; Gibco), 2% (v/v) 33.3 mM glucose (G6152; Sigma-Aldrich, St. Louis, MO), 20 ng per ml Nerve Growth Factor (13290-010; Invitrogen, Carlsbad, CA) and 10 μl/ml Penicillin-Streptomycin-Glutamine (10378-016, Gibco). L-15 medium contains 1.26 mM CaCl_2_. In addition serum contributes about 0.25 to 0.35 mM calcium to the final medium.

### Neuronal cell culture

Fertilized Giriraja-2 chicken eggs were acquired from the Karnataka Veterinary, Animal and Fisheries Sciences University, Bangalore, India. A total of 96 chicken embryos were used in this study. Eggs were incubated at 37 ^∘^C for 8-9 days and dissected in HBSS buffer (14025-092, Gibco) under a stereo microscope to isolate Dorsal Root Ganglia (DRGs). After extraction, DRGs were rinsed in the HBSS buffer lacking Ca^++^ or Mg^++^ (14025-092, Gibco) (14175-095, Gibco), incubated at 37 ^∘^C with 0.5% Trypsin-EDTA (15400-054, Gibco) for 10 min, and then dissociated by gentle pipetting. The cells were seeded on clean, uncoated glass coverslips. Cells were incubated at 37

°C for 96 hr to allow for growth. It is known that by this stage, the membrane associated periodic skeleton is fully developed (Dubey et al., 2020). Before performing ablation experiments, neurons were incubated for 30 min in L-15 medium lacking methylcellulose but containing all the other supplements mentioned above.

### Laser ablation setup and imaging

A home-built laser ablation setup was used to perform the partial ablation experiments. This setup consists of a 355 nm, 25 μJ, 350 ps pulsed laser (PowerChip PNV-M02510-100; Teem Photonics, Meylan, France) coupled to the side port of a Leica TCS SP8 confocal microscope using a custom filter-cube and focused on the sample using 40×/0.75 NA dry phase-contrast objective. A custom-made 90^∘^ rotated filter cube having a UV-reflecting dichroic filter (T387lp-UF3, Chroma) was used to reflect the laser light into the objective. During and after ablation, Phase Contrast images were recorded using a CCD camera (DFC365 FX, Leica) at 15 fps initially and then at 6 fps to capture the initial fast and subsequent slower retraction process of the ablated axons.

### Calcium imaging and chelation

To detect any elevation of calcium levels, cells were preloaded with Fluo-4 AM (F14217, Invitrogen) at 0.5 μM concentration and 20 min of incubation. Chelation of extracellular or intracellular calcium was done using EGTA (03777-10G, Sigma-Aldrich) at 5 mM or BAPTA AM (B6769, Invitrogen) at 10 μM, respectively. The EGTA stock solution was prepared in Millipore deionized water at a concentration of 100 mM. The working concentration of 5 mM was prepared in L-15 medium lacking methylcellulose. The pH of the working solution was adjusted to within

7.2–7.3 by adding either NaOH or HCl and measured using a pH meter. BAPTA was prepared using DMSO as a solvent. The control experiment for Ca^++^ imaging was done using DMSO as a vehicle. Images were acquired using a Leica TCS SP8 confocal system, keeping all the imaging parameters exactly the same before and after ablation.

### Cytoskeletal drugs

To investigate the role of the axonal cytoskeleton in laser ablation-induced retraction, neurons were treated with the following pharmacological agents: 30 *μ*M Blebbistatin (B0560, Sigma-Aldrich) for 20 min to inhibit myosin-II activity, 1 *μ*M Latrunculin-A (L5163, Sigma-Aldrich) for 10 min to depolymerize actin filaments, 16.67 *μ*M Nocodazole (M1404, Sigma-Aldrich) for 30 min to destabilize microtubules, 10 *μ*M Taxol (T7402, Sigma-Aldrich) for 30 min to stabilize microtubules, 5 *μ*M Jasplakinolide (J7473, Thermo Fisher Scientific) for 30 min to stabilize actin filaments. All drugs were dissolved in dimethyl sulfoxide (DMSO) (D84818-50ml, Sigma-Aldrich) with a final DMSO concentration kept below 1% v/v. Control experiments were done with ≤ 1% DMSO.

### Visualization of cytoskeletal dynamics

SPY555-Tubulin dye (SC203, Spirochrome, Switzerland) and SPY650-FastAct dye (SC505, Spirochrome, Switzerland) were used to stain microtubules and actin filaments, respectively. These dyes are specific to these biopolymers and do not label tubulin or actin unless they are in polymerized form. The neurons were incubated with the dye using L-15 media lacking methylcellulose at a concentration of 1:1000 v/v for 30 min to 1 hour before starting the ablation experiments.

### EB3 Transfection and live-cell imaging

To visualize the dynamics of microtubule plus-ends, primary dorsal root ganglia were dissected from 9-day-old chick embryos and transfected with the pCAG-mNeon-EB3 construct via electroporation using a Nepagene electroporator. After electroporation, the cells were plated onto uncoated glass coverslips and maintained in a neuronal growth medium. Following 4 days in vitro (DIV), live-cell imaging was performed using a Leica TCS SP8 confocal microscope in widefield fluorescence mode with a 63 × /1.4 NA oil immersion objective. The mNeon fluorescence was detected using a GFP filter cube (excitation: 450-490 nm, emission: LP 515 nm). Time-lapse sequences were acquired at 5-second intervals to capture the dynamics of EB3-decorated microtubule plus-ends. Kymographs were generated from the time-lapse data using the KymoResliceWide plugin in FIJI/ImageJ software.

### Statistical analysis

Statistical analyses were performed using OriginPro version 2026. Data normality was assessed with the Shapiro-Wilk test. Because at least one group in each dataset failed to meet the normality assumption, non-parametric tests were used for all comparisons. For Figure 3c, normalized Fluo-4 fluorescence intensity values, calculated as I(post)/I(pre), were compared among control, EGTA-treated, and BAPTA-AM treated axons using a Kruskal–Wallis test followed by Dunn’s multiple-comparison test. For Figure 8, actin and microtubule intensity ratios were compared using a two-tailed Mann–Whitney U test. Sample sizes are stated in the corresponding figure captions and Supplementary Table S1. No formal sample size calculation was performed before the experiments. Sample sizes were determined based on successful primary DRG cultures, successful live-cell imaging and laser-ablation experiments, and the availability of analyzable axons for each experimental condition. No statistical test for outliers was performed, and no data points were excluded from the statistical analyses. A p-value of less than 0.05 was considered statistically significant.

## Results

### Partial laser-ablation of axons

Axonal transection studies using lasers are usually performed by locally ablating the axon using a pulsed laser of sufficient pulse energy. Here, we use a method where an axon is ablated with a lower pulse energy than that is required for complete transection so that the mechanical continuity of the cytoskeletal components can be compromised without causing any catastrophic damage to the membrane. In such cases, one can observe two retraction fronts propagating away from the ablation point–one towards the growth-cone and the other towards the soma (see **Figure 1a** and Video 001). Partial ablation events are easily distinguishable from complete transections as (i) a thin membrane connection is visible between the retracting fronts when imaged in phase contrast mode, and (ii) the two retracting ends remain more or less aligned along the initial axis of the axon. This is in contrast with complete transection, where the axonal segments exhibit pronounced buckling and the ends “fly-away” from the initial axonal axis (Video 002). To assess the extent of damage caused by the laser, we performed control experiments using paraformaldehyde-fixed axons. In such axons, no retraction response is observed and the laser induced damage is limited to less than 2 *μ*m (Suppl. Mat., Fig. S1, Video 003).

**Figure 1:**
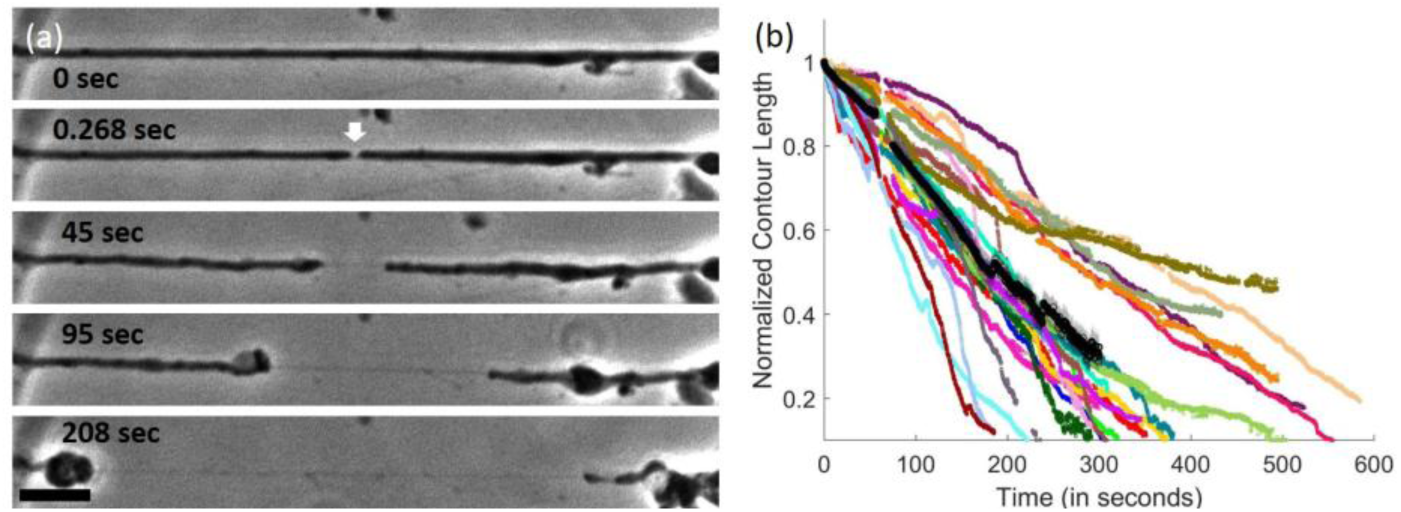
Axonal response to partial laser ablation. (**a**) Sequence of phase contrast images showing the response of an axon to partial laser ablation. The white arrow indicates the point of ablation and the ablation was performed at time t = 0 s. Post ablation, the axon exhibits two retraction fronts–one propagating towards the soma and the other towards the growth-cone. A thin membrane tube spans the region between these two fronts. The scale bar is 10 μm. **(b)** Plots showing the time evolution of the normalized contour lengths of the retracting segments (length of each retracting segment divided by its initial length). Data for individual segments are shown using different colors and the average of all segments is shown using black circles. The average is calculated only up to 300 s to reduce the bias as many retraction events are complete by this time. The gap in the data at around t ≈ 57 s is due to the switching of the recording speed from 15 fps to 6 fps. Shaded region around the average plot represents the standard errors of the mean.

The thinness of the tube between the retracting fronts of partially ablated axons suggests that all cytoskeletal components, along with much of the cytoplasm, retract toward either side post ablation. This will be discussed later with the help of fluorescence images. When cells are grown on untreated coverglass, without any adhesion-promoting treatments, most axons tend to be free of the glass surface except at the soma and the growth-cone ends. In such cases, the two retracting fronts of a partially ablated axon retract continuously until all the material is absorbed into either the soma or the growth cone. This can be seen by analyzing the normalized length of each segment and plotting it as a function of time, as shown in **Figure 1b**. In cases where the distal and proximal side could be clearly identified, no significant difference in retraction speed could be observed (Suppl. Mat., Fig. S2).

### Cytoskeletal perturbations alter axonal retraction

Retraction dynamics following ablation are affected by pharmacological perturbations applied to the cytoskeleton (**Figure 2**). In control conditions, axonal segments show a gradual decrease of their contour lengths over time. Disruption of microtubules with Nocodazole (Noco) leads to a faster and more pronounced retraction, whereas stabilization with Paclitaxel (Taxol) significantly suppresses retraction. In sharp contrast, disruption of actin filaments using Latrunculin A (LatA) decreases the retraction rate significantly and stabilization of these filaments with Jasplakinolide (Jasp) increases the retraction rate. In agreement with actin disruption, inhibition of myosin-II motors using Blebbistatin (Blebbi) decreases the retraction rate by about the same level as LatA. Moreover, consistent with the above results, a combination treatment of Taxol plus LatA results in almost complete inhibition of retraction. The possible mechanisms responsible for these opposing trends shown by the microtubule and actin skeletons will be discussed in a later section.

**Figure 2:**
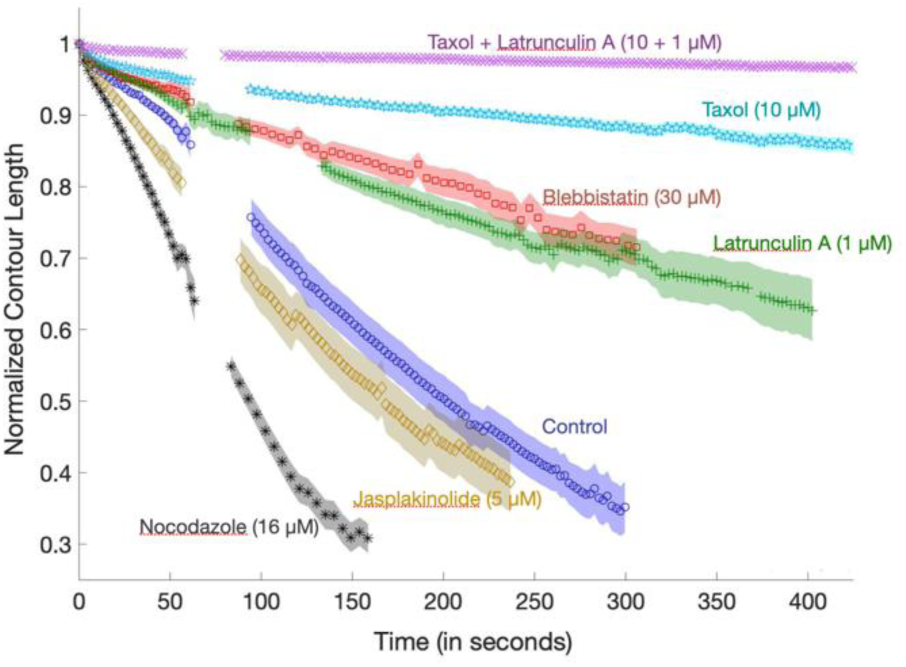
Effect of cytoskeletal perturbations on axonal retraction. Normalized and averaged contour lengths as a function of time following partial laser ablation of axons exposed to different pharmacological agents. Note that the duration of recording for each drug is different due to the difference in retraction rates (see Suppl. Mat., Fig. S3 for full data), and the gap in data is due to the switching of the frame rate of recording. Control axons (n = 24 axonal segments) exhibit sustained retraction over several hundred seconds. Microtubule destabilization using Nocodazole (n = 31) leads to increased retraction speed, whereas microtubule stabilization using Taxol (n = 25) strongly suppresses retraction. In contrast, actin filament destabilization using Latrunculin A (n = 30) decreases retraction speed whereas stabilization using Jasplakinolide (n = 25) increases the rate. Inhibition of myosin-II using Blebbistatin (n = 28) shows a similar effect as actin filament disruption. Combined treatment with Taxol and Latrunculin A (n = 34) almost completely suppresses retraction. Each drug was added 30 min prior to ablation. Shaded regions represent the standard error of the mean.

### Partial laser-ablation causes Ca^++^ elevation in axons

Calcium has well-documented but complex roles in axonal injury and regeneration (Bradke et al., 2012), which motivated us to ask whether it also drives the retraction we observe after partial ablation. In order to explore the possible role of free calcium in driving axonal retraction after partial ablation, we imaged intracellular free Ca^++^ levels using Fluo4-AM dye. As can be seen from ***Figure 3*a**, there is a sudden and significant increase in the intracellular Ca^++^ level when the axon is ablated (Video 004). To identify the mechanisms of calcium elevation, we pretreated cells with either EGTA (5mM), which chelates extracellular Ca^++^, or with the cell permeable intracellular Ca^++^ chelator BAPTA-AM (10*μ*M). Both treatments almost completely suppress Ca^++^ elevation in ablated axons, and example images of an axon treated with EGTA and then ablated is shown in ***Figure 3*b**. The quantification for both treatments is shown in ***Figure 3*c**. These data show that the elevation in Ca^++^ is triggered by entry of extracellular Ca^++^, possibly at the ablation point, which triggers the release of Ca^++^ from internal stores. Measurement of the Fluo4-AM fluorescence intensity shows that the Ca^++^ elevation post-ablation is sudden and reaches peak intensity within a few seconds. This is followed by an exponential decay of free calcium levels as can be seen from the inset of ***Figure 3*c**, with a decay time of 42.7 s. In comparison to this ablation-induced response, mechanical indentation of soma of rat cortical neurons evoked a calcium response in the axon, with a decay time constant of approximately 24.1 s (Gaub et al., 2020). The calcium level seems to decay to a value that is higher than the pre-ablation value but this could be because of the limited observation time in our experiments.

### Chelation of Ca^++^ allows for axonal recovery post ablation

The fact that either EGTA or BAPTA-AM can inhibit calcium elevation suggests that calcium entry from outside is necessary to trigger the internal stores. So, we quantified the post-ablation response of axons maintained in calcium chelated medium. When free Ca^++^ was chelated using EGTA (5 mM), retraction was partial. More remarkably, a subset of axons (10/20 axons) recovered their diameter subsequent to an initial thinning down of their mid-section (see **Figure 4a**). This initial retraction occurred in both the proximal and distal axonal segments, but subsequent recovery happened mostly from the proximal side, as shown by microtubule plus-tip markers discussed later. This is quantified by calculating the contour length of each segment, and the data for the proximal and distal sides are displayed in **Figure 4b** (also see schematic Suppl. Mat., Fig. S4)). A subset of axons retracts partially after ablation and does not recover within the observation time of about 600 s. Example images and population data for these are shown in **Figure 4c,d**, respectively. The retraction velocity is much less for axons in calcium free medium as compared to control axons, as can be seen in **Figure 4d**.

**Figure 3:**
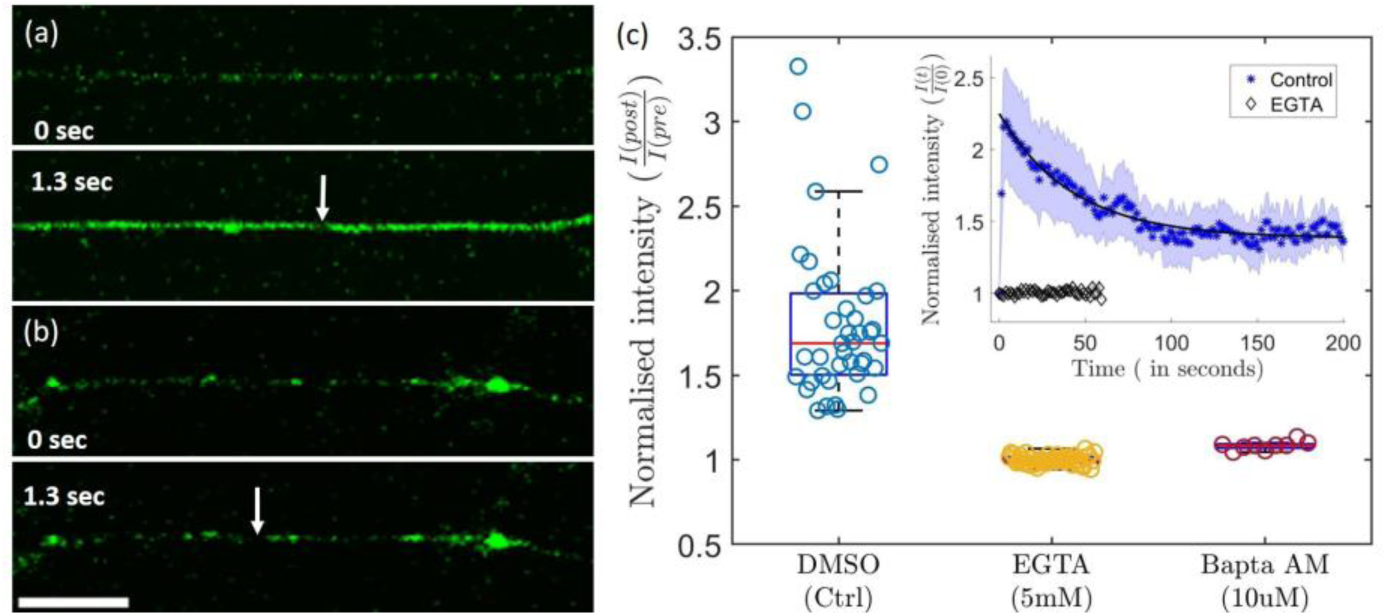
Partial ablation causes Ca^++^ elevation in axons. (**a**) Images of an axon labeled with the Ca^++^ indicator Fluo4-AM taken before ablation (above) and immediately after ablation (below). A sudden increase in fluorescence intensity is seen in every ablated axon. The white arrow marks the point of ablation and the scale bar is 20 μm. **(b)** Images of an axon incubated in medium containing EGTA and ablated as before. No significant increase in fluorescence is seen in this case. **(c)** Data showing the fluorescence intensity normalized with the pre-ablation intensity (I(post)/ I(pre)): for control axons (n = 40), axons grown in medium containing EGTA (n = 47) and for axons treated with BAPTA AM (n = 9). The concentrations used are indicated in the plot. The error bars are standard errors of the mean. **(Inset)** Plots showing the time evolution of intracellular free calcium for control axons and for axons in medium containing EGTA. Ablation was performed at time t = 0 s. The data points are the averages, and the shaded regions are standard errors of the mean for control axons (n = 5) and EGTA-treated axons (n = 7). A sharp increase in Ca^++^ followed by a decay can be seen for the control axon, whereas the EGTA treated axon shows a nearly constant base value of intensity. For control axons, a fit to an exponential function is shown using a solid black line, which yields a Ca^++^ decay time of τ = 42.7 s. Statistical analysis for (c) was performed using a Kruskal–Wallis test followed by Dunn’s multiple-comparison test. The groups differed significantly, H(2) = 76.69, p < 0.0001. Dunn’s post hoc comparisons showed significant differences between control and EGTA-treated axons, p < 0.0001; control and BAPTA-AM-treated axons, p = 0.047. All pairwise comparisons are reported in Supplementary Table S1.

**Figure 4:**
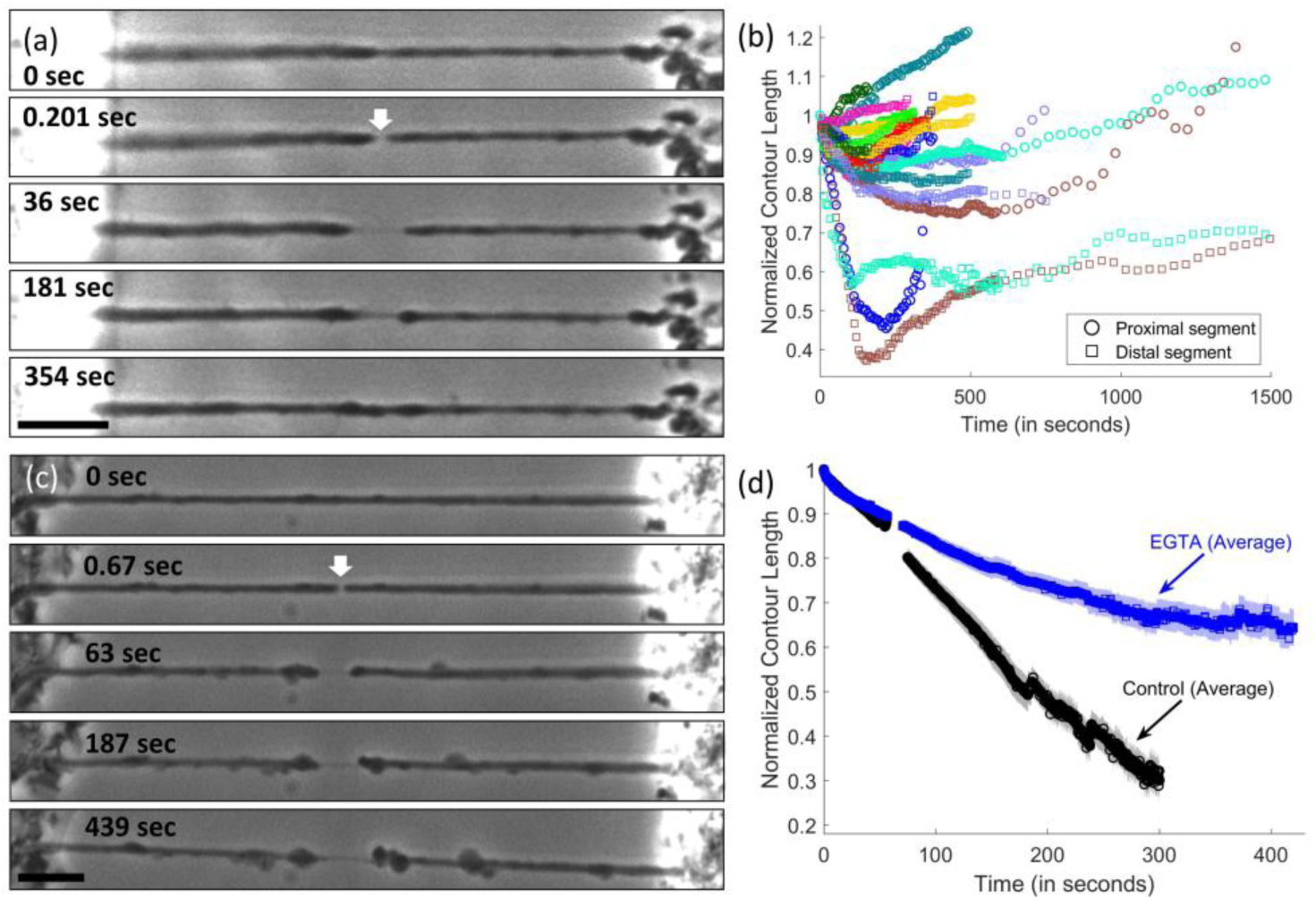
Recovery of damage seen in Ca^++^ chelated medium. (**a**) A sequence of phase contrast images showing the development of a damaged region and subsequent axonal resealing. The cell body is to the left of the images and the white arrow indicates the point of ablation. Both proximal and distal axonal segments initially retract from the ablation point, but recovery mainly occurs from the proximal side. The scale bar is 10 μm. **(b)** The normalized contour length versus time plot for the axonal segments shows the initial retraction and recovery for individual segments. The proximal segment (cell body side) is shown using circles and the distal segment using squares. Segments from the same neuron are shown using the same color. Note that in some cases the normalized length exceeds unity indicating that one segment has grown beyond its initial length at the expense of the other (see schematic Suppl. Mat., Fig. S4). **(c)** Phase contrast images of an axon which was partially ablated in Ca^++^ chelated medium. In this case, the axon has retracted only partially, even after about 400 s. The scale bar is 10 μm. **(d)** Averaged normalized contour length for control axons (same as in **Figure 1b**) and for axons ablated in medium containing 5 mM EGTA. As can be seen from the data, axons maintained in EGTA medium retract only partially, and the retraction velocity is much less than that for control axons. Data for individual EGTA treated axons are shown in Suppl. Mat., Fig. S5. The error bars are standard errors of the mean. The gap in the data at around t ≈ 57 s is due to the switching of the recording speed from 15 fps to 6 fps.

### Axonal microtubules regrow and reseal after ablation in Ca^++^-free medium

The resealing response seen in calcium chelated medium raises the question as to how the axonal cytoskeleton may be responding under such conditions. Microtubules are found in abundance in chick DRG axons, and so it is logical to check their contribution to the retraction and resealing processes. For this, we labeled microtubules using the cell permeable SPY555-Tubulin dye which labels only polymerized tubulin. For control cells grown in medium containing calcium, we see that microtubule intensity recedes in both directions and all the way to the two extremities of partially ablated axons (see **Figure 5a**). This mirrors the retraction response seen in phase contrast images. In some cases, a bulb-like shape is seen at the retracting edge, possibly generated by highly buckled or curled microtubules. Some axons remain straight during the retraction process, while others show some amount of buckling. No detectable fluorescence intensity is seen in the thinned down region, suggesting complete depolymerization of microtubules (see **Figure 5b**), as this region expands with time. A slight increase in intensity is observed at the boundaries of the retracting segments.

**Figure 5:**
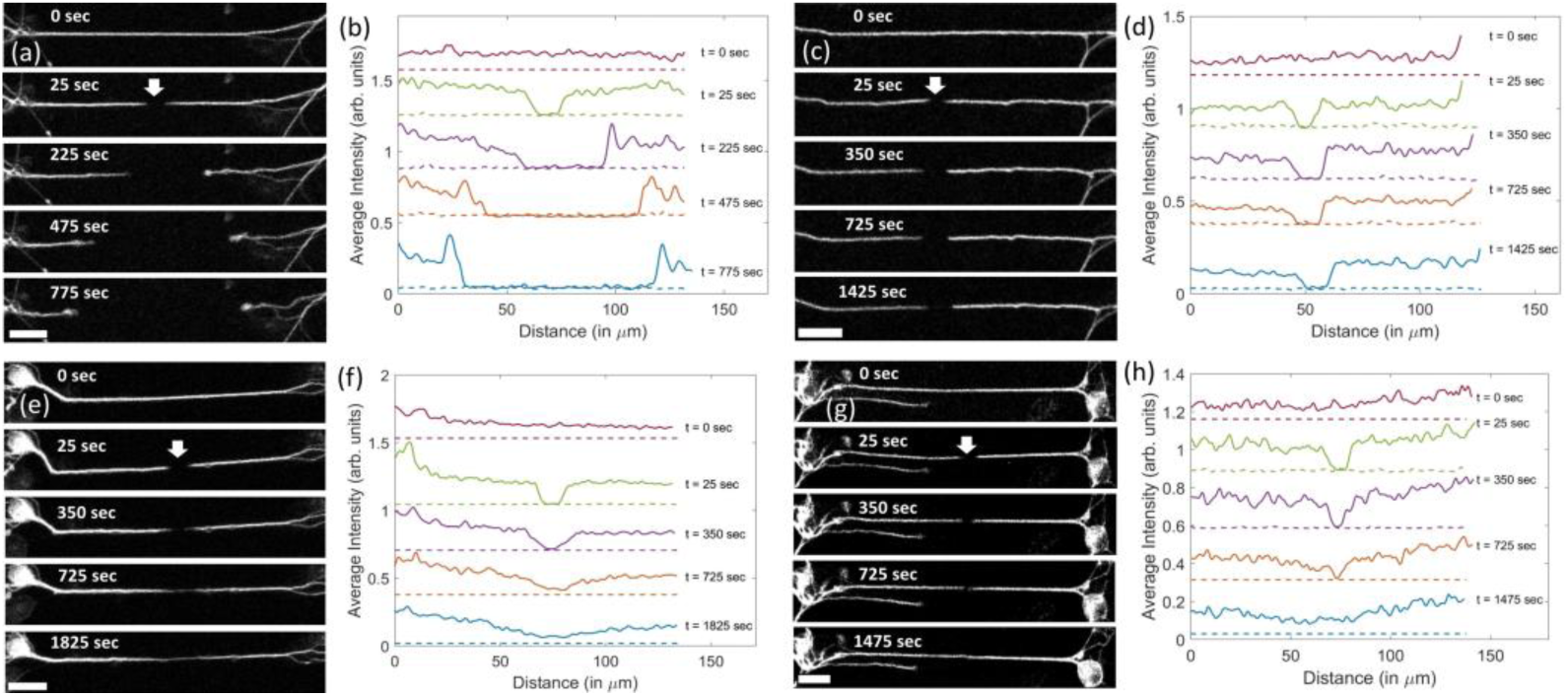
Microtubule dynamics in partially ablated axons. (**a**) Sequence of fluorescence images of an axon labeled with the membrane permeable SPY555-Tubulin dye and partially ablated at the point shown by the white arrow. The dye labels only microtubules and not tubulin dimers. Microtubule intensity retracts along with the retraction front seen in phase contrast images. **(b)** Plots of the microtubule fluorescence intensity corresponding to the images shown in (a). No detectable intensity can be seen in the thinned down mid-section of the axon–the dotted lines indicate background intensity. A slight accumulation of microtubules can be seen at the retraction fronts. **(c)** Fluorescence images showing the microtubule retraction in an ablated axon where calcium in the medium was chelated using 5 mM EGTA. In this case, microtubules retract only partially and then remain static for about 25 min (observation time). **(d)** Plots showing the time evolution of the intensity reduction in the thinned-out regions for the axon shown in (c). **(e,f)** Image sequence and intensity plots for an axon that exhibited only partial recovery of microtubules when ablated in Ca^++^ chelated medium. **(g,h)** Fluorescence images and intensity plots for an axon that showed complete recovery of microtubule intensity subsequent to an initial retraction phase when ablated in Ca^++^ chelated medium. In all cases, recovery of microtubules correlated with recovery of diameter as seen from phase contrast images (see Suppl. Mat., Fig. S6, Video 005). The scale bars are 20 μ m. In all cases, intensity plots were constructed by smoothing the raw intensity data with a Gaussian filter (σ = 1.4) in Matlab and then integrating along the axonal thickness.

Next, we explored how microtubules behave when extracellular Ca^++^ is chelated. Axons of neurons grown in medium containing 5 mM EGTA show different kinds of responses after partial ablation. Some axons showed an initial retraction of microtubule intensity after which the intensity evolution stops and remains stable for the rest of the observation time of about 25 min (see **Figure 5c,d**). Comparison of phase contrast and fluorescence images shows that the regions devoid of microtubules appear thin, while the parts where intensity is unaffected have normal caliber. In other cases, the intensity initially recedes, then stops receding, starts to recover, and eventually recovers either partially (**Figure 5e,f**) or completely (**Figure 5g,h**). The time evolutions of microtubule intensity for different axons mirror the observations of diameter measured for the same axons. This suggests that the microtubule content in axons is directly correlated with axonal diameter (see Suppl. Mat., Fig. S6).

Next, we explored how the microtubule recovery happens in the subset of axons that showed recovery when maintained in calcium chelated medium. For this, we transfected neurons with mNeon tagged EB3 to visualize the plus-end dynamics of growing microtubules (Stepanova et al., 2003). Note that in DRG axons microtubules are arranged with a nearly perfect polar order with their plus tip pointed towards the growth cone end. This can be seen in control axons where EB3 comets travel towards the growth cone end with only very few comets in the opposite direction (see Suppl. Mat., Fig. S7). As can be seen from **Figure 6** (see Suppl. Mat., Fig. S8 for more examples), during the post ablation recovery phase, this directional bias was maintained, with a higher comet density near the recovering proximal ends. This is consistent with the predominantly proximal recovery observed in **Figure 4b**. This suggests that recovery may be happening either through regrowth of existing microtubule fragments with pre-established polarity or biased *de novo* nucleation and growth. If it is the latter case, the biasing mechanism is unclear. One possibility is that augmin mediated, templated nucleation propagates the original plus-end-out polarity, a mechanism shown to orient axonal microtubules (Sánchez-Huertas et al., 2016).

**Figure 6:**
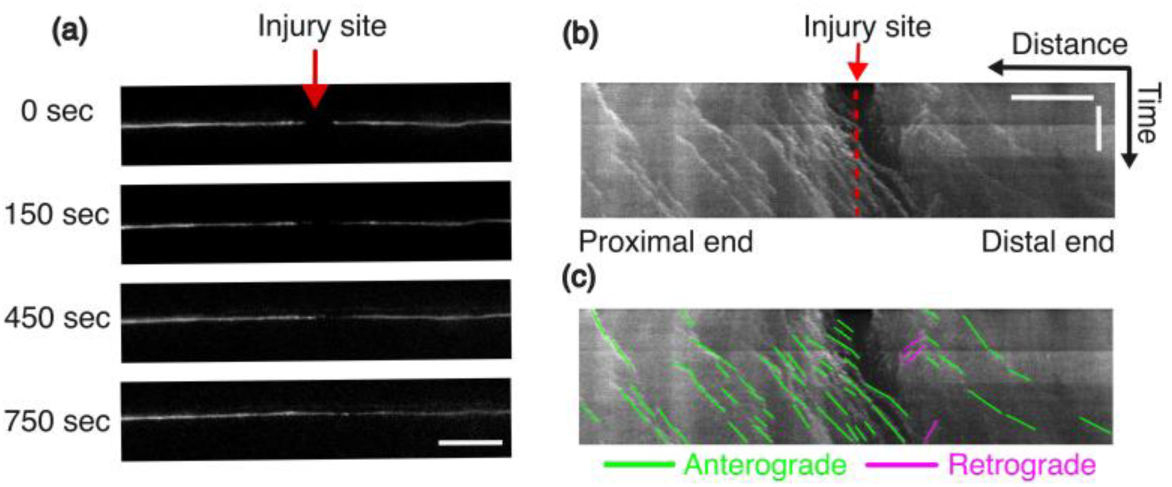
Visualization of microtubule regrowth in Ca^++^ chelated media. (**a**) Selected frames from a time-lapse movie showing EB3 fluorescence intensity during axonal recovery following partial laser ablation (see Suppl. Mat., Figs. S7, S8 and Video 006). The injury site is indicated by a red arrow. The cell body is to the left of the images. Scale bar: 10 μm. **(b)** A kymograph generated along the axon showing EB3 dynamics during the recovery phase. A red arrow and a dotted red line mark the injury site. The horizontal axis represents distance along the axon (scale bar: 10 μm), and the vertical axis represents time (scale bar: 5 min). **(c)** The same kymograph as in (b), with overlaid traces highlighting EB3 comet trajectories. Green lines represent anterograde growth, while magenta lines indicate retrograde ones. Representative trajectories are shown for illustration only. The proximal side of the injury demonstrates a higher concentration of tracks, which aligns with site-specific microtubule regrowth.

### Actin filaments too recover in calcium-chelated medium

Compared to microtubule organization, axonal actin filaments form more diverse structures. Mature axons contain a membrane associated periodic skeleton (Xu et al., 2013) as well as other highly heterogeneous cortical actin structures like actin hot spots and trails (Leterrier et al., 2017). Although less abundant, these structures can contribute significantly to axonal mechanics and retraction dynamics (Costa et al., 2020; Datar et al., 2019; Dubey et al., 2020; Fan et al., 2019; Mutalik et al., 2018). Note that the membrane associated actin-spectrin periodic skeleton is fully developed in the four days in culture chick DRG neurons we use (Dubey et al., 2020). To study the damage and possible recovery of actin filaments, we imaged actin using the cell permeable SPY650-FastAct dye which labels only polymerized actin.

In control experiments, where Ca^++^ is present in the medium, actin fluorescence is seen to recede completely as in the case of microtubules (see **Figure 7a**). Very prominent actin peaks can be observed at the retracting fronts on either side, suggesting significant accumulation of filaments at these locations (**Figure 7b**). No actin filaments could be detected in the thinned mid-section within the observation time of 30 min. At first sight, this complete loss of actin appears to sit uneasily with the pharmacological data of **Figure 2**, where destabilizing actin slowed retraction; we address this apparent discrepancy in the Discussion. When extracellular Ca^++^ is depleted using EGTA, we observe recovery of actin filaments in the thinned-out region. Different behaviors are observed in such axons. Unlike microtubules, actin filaments recover in all axons maintained in Ca^++^ chelated medium–irrespective of whether the axonal diameter recovers or not. Unlike in the case of microtubules, after an initial decrease and subsequent recovery, the intensity often overshoots significantly–exceeding its pre-ablation value as can be seen in **Figure 7c,d** and is quantified in ***Figure 8***. This excess actin eventually redistributes. The recovery pattern too is different from what was observed for microtubules under similar conditions. No clearly demarcated growing fronts are observable in the case of actin filaments. Instead, actin recovery is distributed all along the axon and can be heterogeneous, as can be seen in **Figure 7e,f**. Another notable difference, when compared to microtubule recovery, is in the correlation between recovery of axonal diameter and actin filaments. In **Figure 7g** we show an example of an axon which has fully recovered its diameter as well as actin filament intensity. However, in **Figure 7h**, one can see a case where actin intensity has recovered fully, but the diameter failed to recover within the same time span.

**Figure 7:**
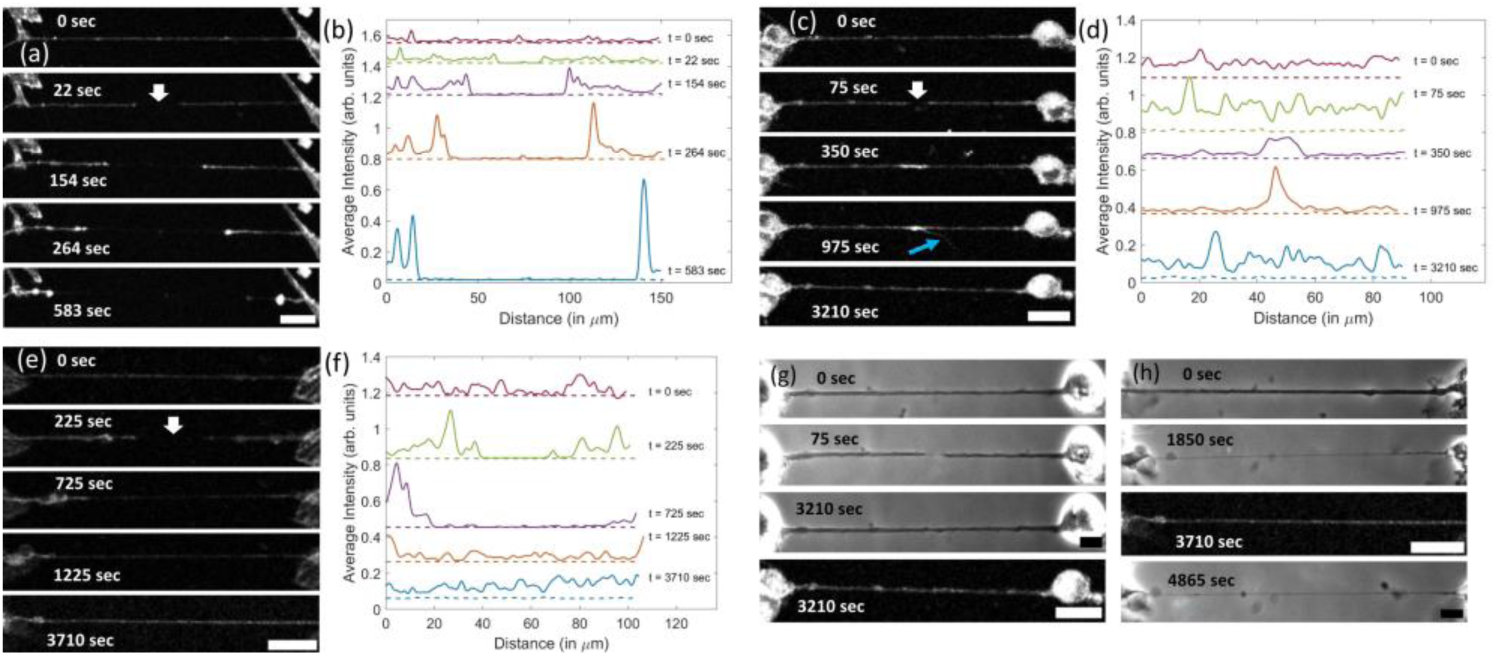
Recovery of actin filaments post ablation in Ca^++^ chelated medium. (**a**) Sequence of fluorescence images of a control axon which was labeled using SPY650-FastAct dye which labels only polymerized actin and was partially ablated. The white arrow indicates the point of ablation. Post-ablation, a thin tube spans the region between the retracting edges (not visible in the fluorescence images). Actin filaments tend to accumulate at the retracting edges of each axonal segment. **(b)** Intensity plots showing the time evolution of actin fluorescence. Hardly any actin intensity is seen in the thinned-out regions. The dotted lines represent the zero intensity level. **(c)** When partial ablation is performed in calcium chelated medium, actin intensity in the thin region initially decreases (t = 75 s), then recovers, often overshoots (t = 350 s), and finally redistributes (t = 3210 s) (Video 007). The blue arrow indicates filopodia-like structures that form at regions with enhanced actin intensity. **(d)** Quantification of the actin fluorescence post ablation and during recovery for the images shown in (c). The overshoot is clearly visible at t = 350 & 975 s. **(e,f)** Another example of an axon maintained in calcium chelated medium. In this case, retraction is much more extensive, making it easier to see how actin filaments recover in the region between the retraction fronts. Unlike microtubules, recovery of actin intensity is seen all along the thinned-out segment (t = 725 & 1225 s). A slight overshoot can be seen in this case too (t = 3710 s). The corresponding intensity plots are shown in (f). To obtain all the intensity plots, the raw intensity data have been smoothed with a Gaussian filter (σ = 1.4) using Matlab. The intensity is then integrated across the axonal thickness. **(g,h)** While all axons showed recovery of actin filament intensity, the recovery of actin filaments may or may not be correlated with the recovery of axonal diameter. In (g), we show the example of an axon that exhibited diameter recovery as well as actin recovery. Whereas, in (h) one can see that there is no recovery in diameter even though actin intensity has recovered all along the axon. A phase contrast image at a much later time point shows that the diameter remains small for long times. The scale bars in the fluorescence and phase contrast images are 20 μm and 10 μm, respectively.

**Figure 8:**
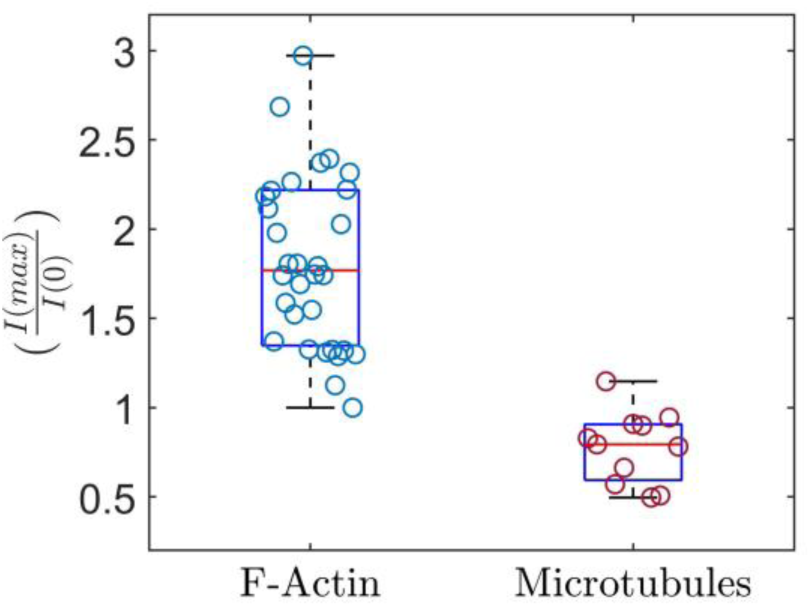
Comparison of post-ablation intensity of microtubules and actin filaments. The data show the ratio of maximum post-ablation intensity per unit axonal length I(max) to the pre-ablation intensity per unit length I(0), measured around the ablation point where actin filaments and microtubule initially retracted and later reformed. Significant overshoot in intensity is observed in the case of actin, even when there is no diameter recovery. In the microtubule case, some axons showed near complete recovery whereas others did not fully recover within the observation time (note that recovery time depends on the extent of initial retraction as well). The error bars represent values within 1.5 times the interquartile range from the first and third quartiles, illustrating the range of non-outlier data. Sample sizes were n = 32 actin-labeled axons and n = 11 microtubule-labeled axons. Statistical analysis was performed using a two-tailed Mann–Whitney U test (Supplementary Table S1). Actin and microtubule intensity ratios differed significantly, U = 350, p <0.0001.

## Discussion

### Cyto-mechanics of axonal retraction after partial laser-ablation

When control axons are partially ablated, a thinned down membrane segment develops at the ablation point and expands in either direction, indicating cytoskeletal atrophy (**Figure 1**). The pharmacological perturbation data show that axonal stability post-ablation is dictated by a balance of acto-myosin contractility and microtubule stability (**Figure 2**). Either disruption of actin filaments or inhibition of mysoin-II decreases the retraction speed by comparable levels; and stabilization of actin filaments enhances retraction. In contrast, disruption of microtubules enhances retraction, and stabilization of tubules results in drastically reduced retraction. These observations suggest that the retraction is driven at least partially by acto-myosin contractility, with microtubules resisting this process (P. Baas & Ahmad, 2001; Gallo, 2004; Mutalik et al., 2018). The exact mechanism by which the actin-spectrin membrane associated periodic scaffold generates compressive stresses is yet to be fully understood (Mikhaylova et al., 2020). Microtubules could release compressive stresses generated by myosin-II by either sliding with respect to each other or by en masse depolymerization. As discussed below, imaging of microtubules in retracting axons reveals the possible mechanism.

The pharmacological finding that actin and myosin-II promote retraction, even though actin fluorescence recedes from the injury zone (**Figure 7**), can be reconciled spatially: acto-myosin may generate contractile stress in the intact segments flanking the injury while disassembling at the advancing retraction front itself, so that the filaments producing force are not those being lost. This interpretation remains to be tested directly. Pharmacological stabilization, on the other hand, increases the retraction speed only slightly (**Figure 2)**. This may be either because myosin density is not significantly altered by this treatment or stabilized filaments alter myosin activity, or because Jasplakinolide increases actin density only slightly.

### Ca^++^ regulates cytoskeletal remodeling in ablated axons

In the partial ablation experiments, we also observe that the action of the laser triggers a transient elevation of cytosolic Ca^++^ in the entire axon within a few seconds (***Figure 3*a**), and this is mediated by extracellular calcium entry and subsequent release from stores. The atrophy occurs via progressive loss of cytoskeletal structures in either direction from the injury site. This process can be delayed, arrested, or even reversed when extracellular calcium is chelated.

We observe that Ca^++^ elevation renders axonal microtubules more susceptible to laser-induced degeneration and complete recovery of microtubules can occur when extracellular Ca^++^ is chelated. Hence, it is likely that depolymerization is driven at least partly by calcium induced mechanisms. Calcium is known to affect microtubule catastrophe frequency in in vitro experiments (O’Brien et al., 1997; Weisenberg, 1972; Weisenberg & Deery, 1981), and in cells, calpain activation by calcium is known to degrade microtubules (Büki & Povlishock, 2006; Ziv & Spira, 1995). Beyond calpain, the Ca^++^ transient may contribute to microtubule destabilization through calcium-responsive regulators. EF-hand Ca^++^ –binding proteins such as calmodulin and S100 can modulate brain microtubule assembly, while Ca^++^/calmodulin-dependent phosphorylation of neuronal MAPs, including MAP2 and tau, can reduce their ability to support microtubule assembly (Baudier et al., 1982; Marcum et al., 1978; Yamamoto et al., 1985). In parallel, active depolymerizers may amplify microtubule loss from newly exposed ends. Kinesin-13 proteins such as MCAK/KIF2C directly destabilize microtubule ends, whereas KIF2A regulates microtubule dynamics in neurons; kinesin-8 proteins represent another class of microtubule-dynamics regulators (Desai et al., 1999; Homma et al., 2003; Walczak et al., 2013). These mechanisms were not tested in our system and are therefore presented as possible contributors to the propagating depolymerization front.

In our partially ablated axons, intracellular Ca^++^ level peaks within seconds and then decays exponentially with a decay time of 42.7 s (***Figure 3*c**, inset). Axonal retraction, however, continues over a few hundred seconds or longer (**Figure 2**). These observations suggest a two-stage process where initial destabilization is driven by Ca^++^ mediated processes and once sufficiently destabilized, the cytoskeleton is prone to continued atrophy. Taxol stabilized microtubules show reduced proteolytic fragmentation (Billger et al., 1988) as well as reduced catastrophe frequency (Jordan & Wilson, 2004; Schiff et al., 1979), consistent with our observation. Actin filaments too are affected by excess Ca^++^, as chelation results in recovery of these filaments throughout the axon. It has been shown recently that the actin-spectrin periodic scaffold remodels under elevated Ca^++^ conditions via multiple enzymatic processes (Heller et al., 2025a). In addition, Ca^++^ can enhance acto-myosin contractility through the calmodulin–MLCK pathway: elevated Ca^++^ binds calmodulin, which activates myosin light-chain kinase (MLCK); MLCK in turn phosphorylates the regulatory light chain of non-muscle myosin-II, promoting contractility (Salbreux et al., 2007).

Calcium is recognized as the primary signal of axonal injury, and its consequences are double-edged. In vertebrate axons imaged in vivo, the Ca^++^ rise evoked by transection unfolds in two phases—a local rise at the moment of injury and a later, spreading wave that precedes fragmentation (Vargas et al., 2015). When large or sustained, this Ca^++^ is destructive: it drives calpain-mediated proteolysis of the cytoskeleton and, through mitochondrial Ca^++^ overload, bioenergetic failure and oxidative stress, engaging the programmed axon-death (Wallerian) pathway (Conforti et al., 2014; Villegas et al., 2014).

It has been shown in literature that the same transient is, however, required for repair of fully transected axons. After complete transection, resealing of the cut end is Ca^++^-dependent—the local Ca^++^ rise triggering vesicle fusion that rebuilds the membrane barrier (Mencel & Bittner, 2023). Extracellular Ca^++^ is likewise required to reorganize the injured end into a new growth cone (Bradke et al., 2012). And the injury-evoked signal travels back to the cell body to switch on the neuron’s growth program (Rishal & Fainzilber, 2014). Whether Ca^++^ harms or heals therefore depends on its magnitude, duration and spatial reach: a brief, locally confined transient favors sealing and recovery, whereas a large or spreading elevation commits the axon to destruction (Rishal & Fainzilber, 2014; Vargas et al., 2015).

The partial ablation experiments reported allow us to quantitatively interrogate calcium mediated effects on the axonal shaft, without interference from membrane resealing and growth-cone regeneration processes. In this scenario, chelating extracellular Ca^++^ mitigates axonal atrophy. Ca^++^ may contribute to atrophy via enhanced acto-myosin contractility, as discussed above, as well as by triggering the collapse of cytoskeletal components. The latter effect is discussed in more detail below.

### Microtubules depolymerize *en masse* in the form of a retracting front

Fluorescence imaging shows that the parallel microtubule array or bundle within the axon depolymerizes progressively from the ablation point forming a receding front. Actin filaments too disappear from the thinned out mid-section but show very significant accumulation near the retraction fronts. These cytoskeletal changes are shown schematically in **Figure 9a**. The pattern of microtubule disruption is noteworthy, as it is known that axons contain labile and stable fractions of microtubules–the stable fraction remains unaffected for long periods under the action of microtubule disrupting drugs like Nocodazole (P. W. Baas et al., 2016; Datar et al., 2019). When treated with Nocodazole, axonal microtubule bundles thin down radially, but leaves a stable core all along the axon (Datar et al., 2019). In partially ablated axons, however, we see an expanding region of complete depolymerization, with the rest of the bundle remaining intact. These observations raise the question as to what causes a progressive decay of the microtubule array in the form of a receding front.

**Figure 9:**
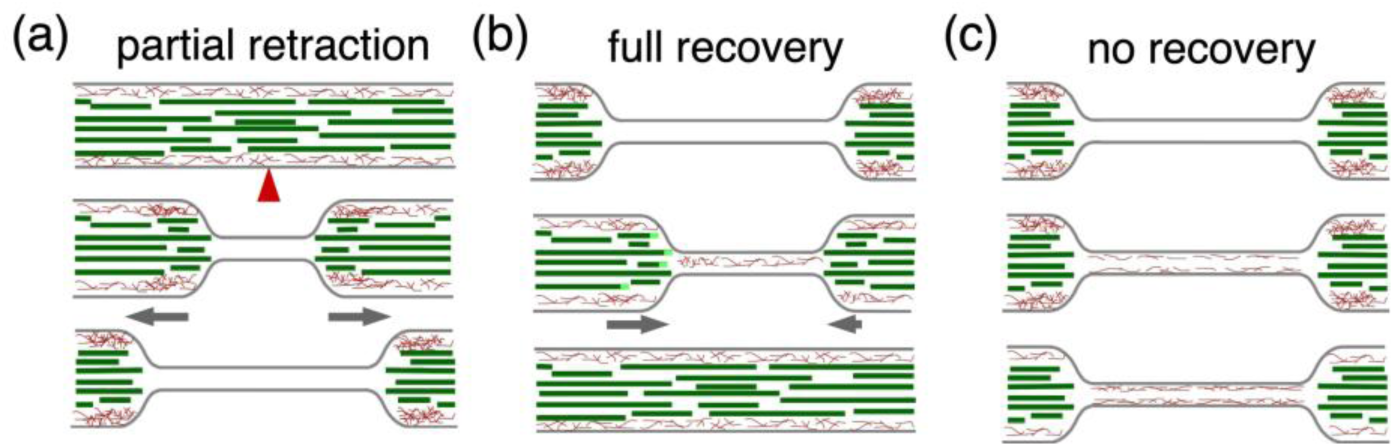
Schematic of cytoskeletal retraction and recovery in Ca^++^ free medium. (**a**) Diagrams showing the receding of microtubules (green) and actin filaments (red) post ablation. The red arrowhead indicates the point of ablation. Although 4 DIV chick DRG axons have well formed periodic actin rings (Dubey et al., 2020), we do not know if the rings reappear when actin filaments recover. For this reason we have not shown actin rings in the schematics. **(b)** Schematic of a case where microtubules and actin filaments recover in calcium chelated medium. Regrowing microtubule tips are marked in light green, and are seen mostly in the proximal segment. **(c)** A case where only actin filaments reform in the thin section. In such cases, there is no recovery of diameter.

Here, we speculate that collective effects arising from microtubule-microtubule interactions may be responsible for the observed dynamics. When microtubules across an axonal cross-section are cut by the laser, these microtubules are left with destabilized ends containing GDP-tubulin and they depolymerize, as shown schematically in ***Figure 10***. The collapse of these microtubules leads to a reduction in the number of neighbors for overlapping microtubules which were not directly affected by the laser, making them unstable. This process continues as depicted in ***Figure 10***. The dependence of microtubule stability on the presence or absence of neighbors may arise for the following reasons. (i) Microtubules may be stabilized by direct Microtubule Associated Proteins (MAPs) based interaction with neighbors. (ii) Loss of neighbors may open up gaps in the array and hence easy access to microtubule severing enzymes like spastin and/or katanin. This may also apply to large proteolytic enzymes like calpains and other Ca^++^ mediated signaling mentioned earlier. Note that even though calcium elevation happens throughout the axon within seconds, microtubule depolymerization proceeds at a much slower rate as a front. (iii) The tubulin post-translational modification (PTM) state of microtubules is altered when PTM modifying enzymes gain access when gaps open up (Janke & Magiera, 2020). This number of nearest neighbor dependent stability hypotheses (NNN-dependent microtubule stability) may account for the catastrophic failure of the microtubule bundle through a cascading effect, as is illustrated in ***Figure 10***.

**Figure 10:**
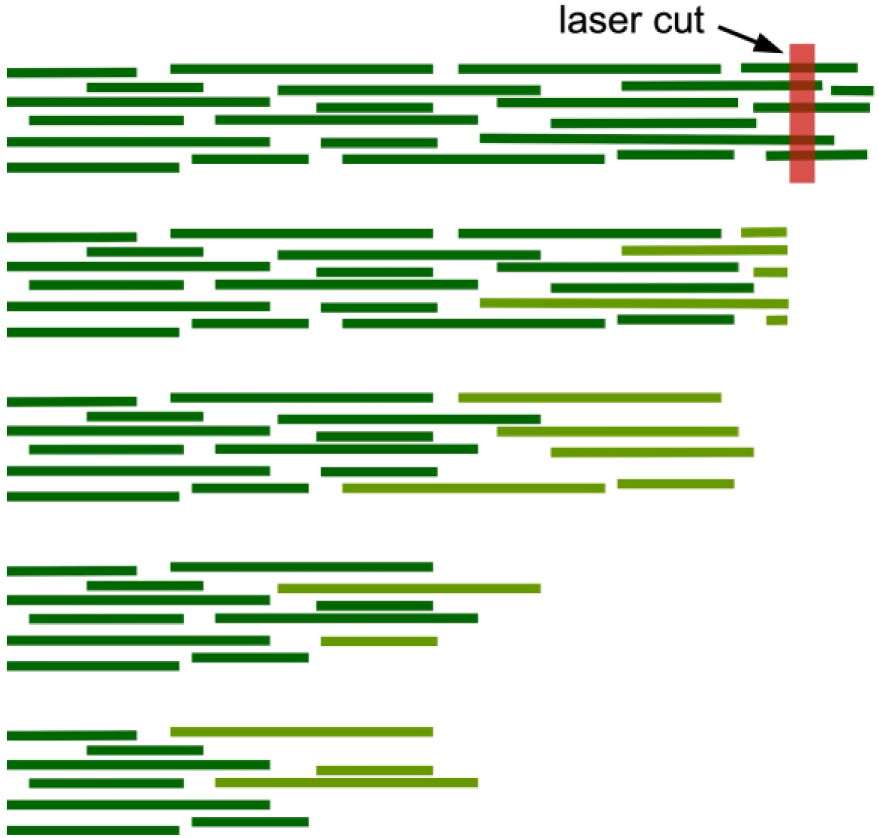
Proposed NNN-dependent microtubule bundle instability mechanism. Schematic showing a possible mechanism for the loss of microtubules within the bundle in the form of a retracting front. Microtubules with compromised stability are shown in light green. Microtubule-associated proteins that interlink microtubules are not shown. Microtubules that are directly cut by the laser become unstable and depolymerize. This leads to a reduction in the number of neighbors for adjacent microtubules, making them unstable–either due to MAP unbinding or due to severing enzymes like spastin and katanin gaining access to loosely packed tubules. This causes a cascade effect leading to a depolymerizing front.

### Microtubules dictate axonal caliber

Remarkably, a fraction of axons shows recovery of normal caliber when calcium is chelated. Imaging of microtubules shows that this requires complete recovery of microtubules (see **Figure 5g,h** and the schematic **Figure 9b**). Furthermore, imaging of EB3 comets shows that microtubule regrowth happens predominantly from the proximal segment. The preservation of microtubule polarity during regrowth suggests that microtubules recover by elongation of existing fragments rather than through *de novo* nucleation. In addition, augmin-mediated branching from existing filaments could also contribute to microtubule recovery (Nguyen et al., 2014; Sánchez-Huertas et al., 2016; Yau et al., 2016). Caliber is classically attributed to neurofilaments, yet in axons that contain few or no neurofilaments—as in many invertebrates, notably arthropods—caliber is dictated by the microtubule array (Prokop, 2020). The relative contribution of neurofilaments in the chick DRG axons used here remains to be established.

Actin filaments too recover after the initial disruption when extra-cellular calcium is chelated. Unlike in the case of microtubules, the reformation of these filaments occurs all along the thinned regions and in several cases exceeds the initial levels (see **Figure 7e,f**). This recovery pattern is reminiscent of the recovery of actin filaments in cellular blebs, where fresh nucleation of filaments occurs from a free membrane soon after the expansion of a bleb, mediated by membrane-bound actin nucleator proteins (Charras et al., 2006). Recovery of actin filaments occurs throughout the entire thinned down segment of all axons maintained in calcium chelated medium. This is in contrast to the formation of a polymerization front in the case of microtubules. Moreover, actin filament recovery alone is not enough for the recovery of axonal caliber (see **Figure 7h** and the schematic **Figure 9c**).

## Conclusion

Axonal degeneration after injury is often regarded as an inexorable consequence of cytoskeletal breakdown. By separating cytoskeletal damage from membrane rupture — using partial laser ablation that spares the plasma membrane — we show that it is a calcium-gated and mechanically driven process that can be mitigated or arrested in multiple ways and, in some cases, reversed. This addresses a gap left by conventional transection models, in which extensive membrane disruption, cytoskeletal loss, and calcium influx occur together and cannot be disentangled, and in which recovery is through growth-cone-mediated regeneration. Through the experiments reported here, we show that the fate of an injured axon is set by coupled factors--the mechanical balance between acto-myosin contractility and microtubule stability and the injury-evoked rise in intracellular free Ca^++^, which drives cytoskeletal disassembly.

Several questions follow. The concentration thresholds at which microtubules become vulnerable to injury-induced Ca^++^ remain to be quantified. If and how neurofilaments recover after partial laser ablation, possibly through microtubule coupled transport (Uchida et al., 2016), also needs to be explored. The nearest-neighbor microtubule stability hypothesis we propose to account for the “burning-fuse” depolymerization front also needs to be tested. More broadly, the demonstration that a partially injured axon can be rescued and rebuilt suggests that combining calcium-targeted intervention with microtubule stabilization and local reduction in myosin-II mediated contractility is a rational route to limiting secondary damage in stretch and crush injuries to nerves. Thus, we believe that these studies open up new avenues in investigating the stability of axonal cytoskeleton and in pharmacological management of axonal injuries.

## Author Contributions

A.M. and P.A.P. designed the research. A.M. and P.J. performed research and analysis. M.A.A. and A.M. developed image analysis codes, A.M., P.J. and P.A.P. interpreted results and wrote the article.

## Declaration

The authors declare no competing interests.

## Acknowledgments

The authors thank the lab of Aurnab Ghose for sharing the pCAG-mNeon-EB3 construct. We acknowledge support through The Wellcome Trust DBT India Alliance (grant IA/TSG/20/1/600137) and through the Indo-Portugal grant, DBT, Gov. of India (DST/INT/Portugal/P-04 /2022).

