## Supplementary material for "Calcium-dependent cytoskeletal collapse and recovery of axons after partial laser ablation": Supplimentry Material

### Supplementary Information for: “Calcium-dependent cytoskeletal collapse and recovery of axons after partial laser ablation.”

Ashish Mishra[1], Pooja Joshi[1], Md Arsalan Ashraf[1], and Pramod Pullarkat*[1]

[1]Raman Research Institute, Bengaluru 560080, India

### Supplementary Material

#### Supplementary figures

##### Ejected length analysis of paraformaldehyde-fixed axons after laser ablation.

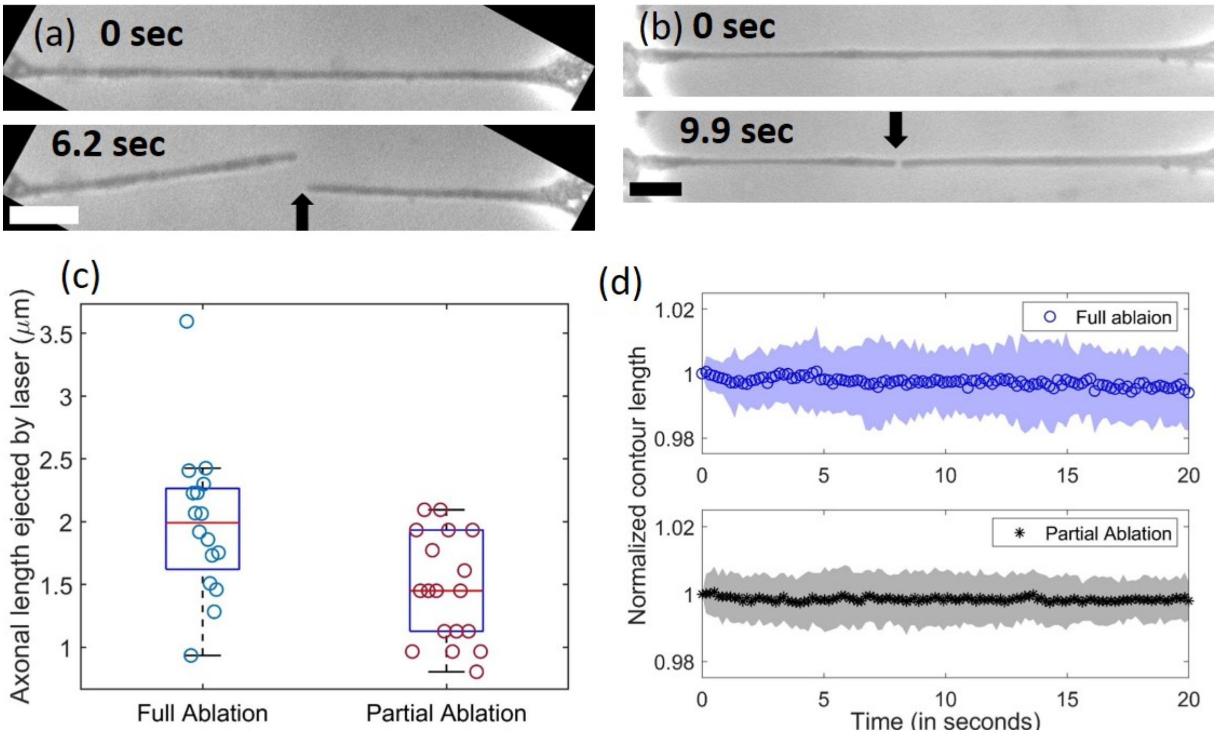


**Fig. S1:** (a) Phase contrast images of a paraformaldehyde (PFA)-fixed axon before and after ablation. The axon has lost less than 2 $\mu$m of its length during ablation and remain stable thereafter. (b) A similar experiment performed at a lower laser power so that the membrane remain intact. In this case the two segments remain aligned due to the membrane bridge between them. (c) Boxplot of axonal lengths ejected by laser during either complete ablation (n = 16) or partial ablation (n = 18) of paraformaldehyde (PFA)-fixed axons. The ejected length was calculated by measuring the stable gap created after injury. (d) Plots of the normalized contour lengths of post-ablation axonal segments (same axons as in (c)). This measurement helps quantify how much of the axon is physically damaged by the action of the laser alone, with minimal effect from cytoskeletal depolymerization (also see Suppli, Video 003). The scale bar is 10 $\mu$m.

**Retraction dynamics of individual axonal segments.**

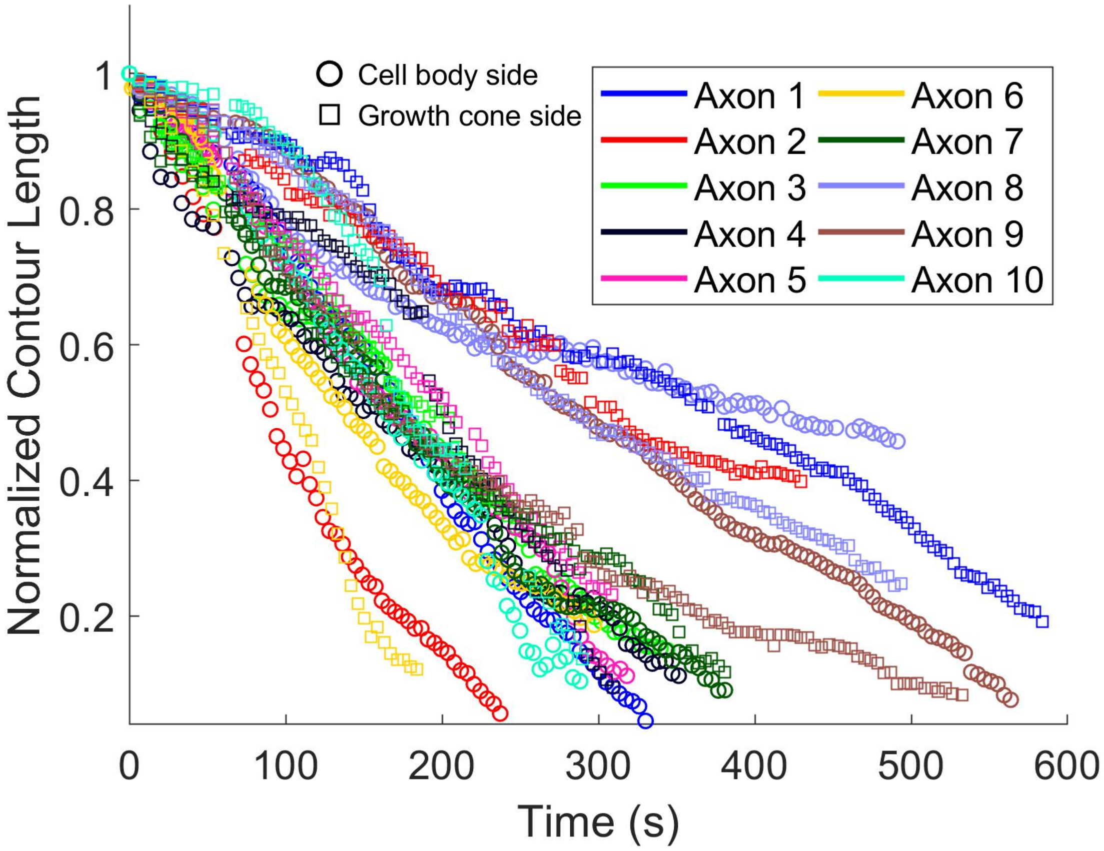


**Fig. S2:** Normalized contour length is plotted as a function of time for individual axons following partial laser ablation under control conditions. Each colour represents a different axon, while circular and square markers denote the cell body side and growth cone side of the same axon, respectively. After partial ablation, both ends of the axon retract over time. While retraction dynamics vary across axons, the behavior of the cell body and growth cone segments is largely similar. Data points are down-sampled for clarity.

#### Retraction behavior of axonal treated with various cytoskeleton perturbing drugs.

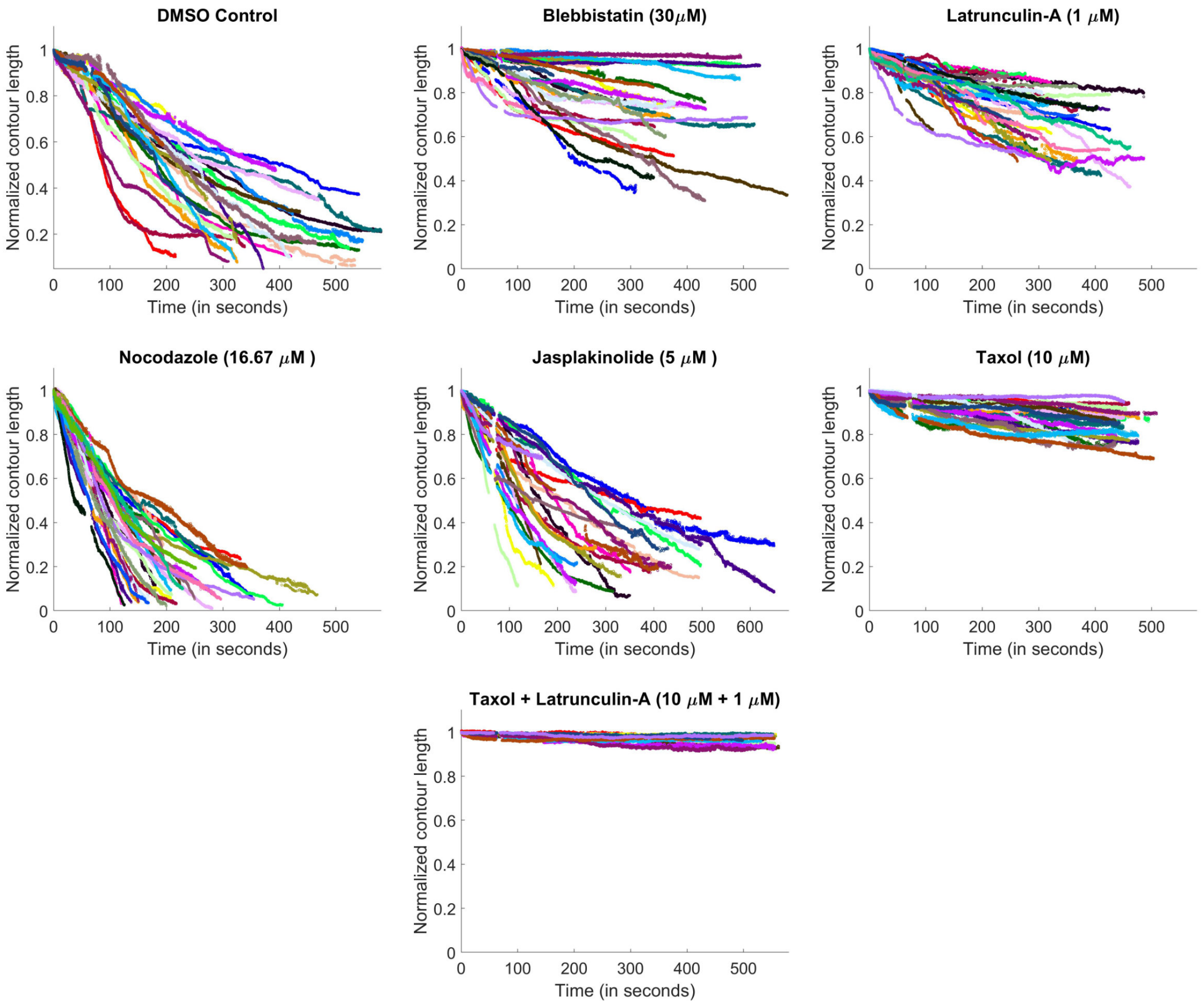


**Fig. S3:** Segment retraction responses observed for axons which were pre-treated with various cytoskeletal drugs. The drug details are indicated on each plot.

**Schematic of axonal segment recovery beyond the ablation point.**

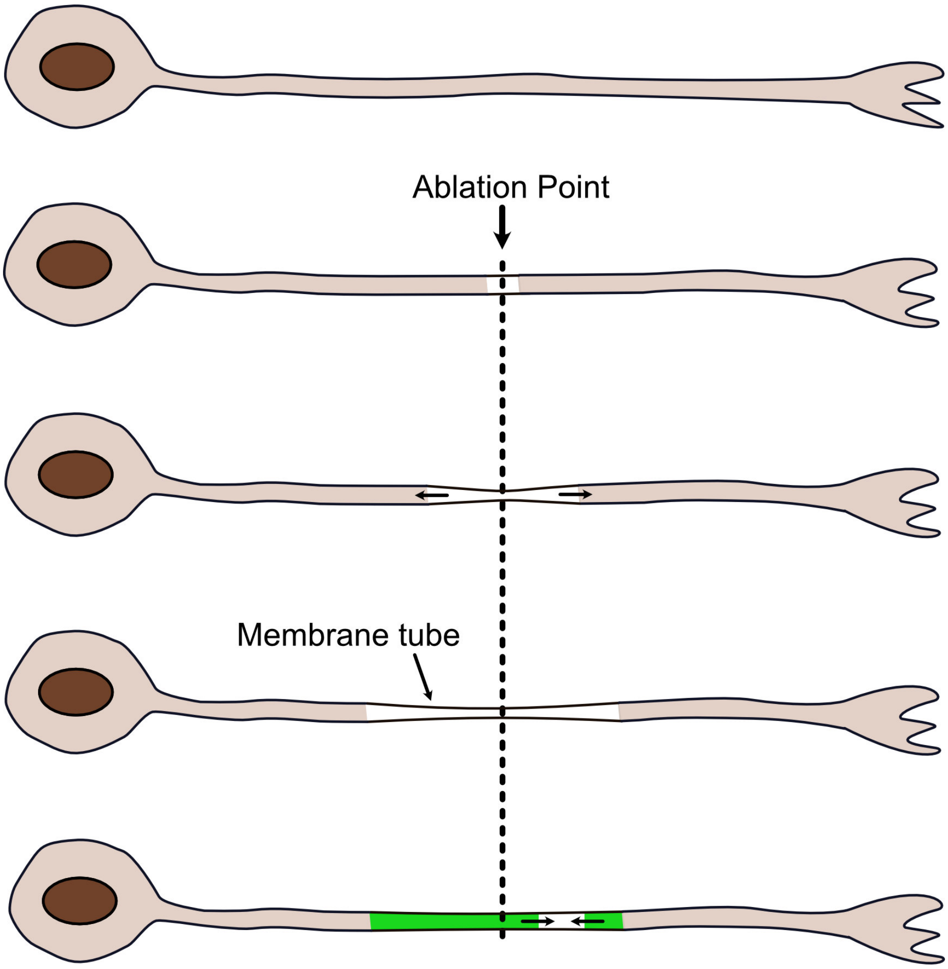


**Fig. S4:** Schematics showing the recovery and resealing of a partially ablated axon. In this case, the regrowth (colored green) is predominantly from the proximal side and hence this segment overshoots the ablation point. The quantitative data is shown in the main text.

**Retraction plots of EGTA-treated axonal segments after partial laser ablation.**

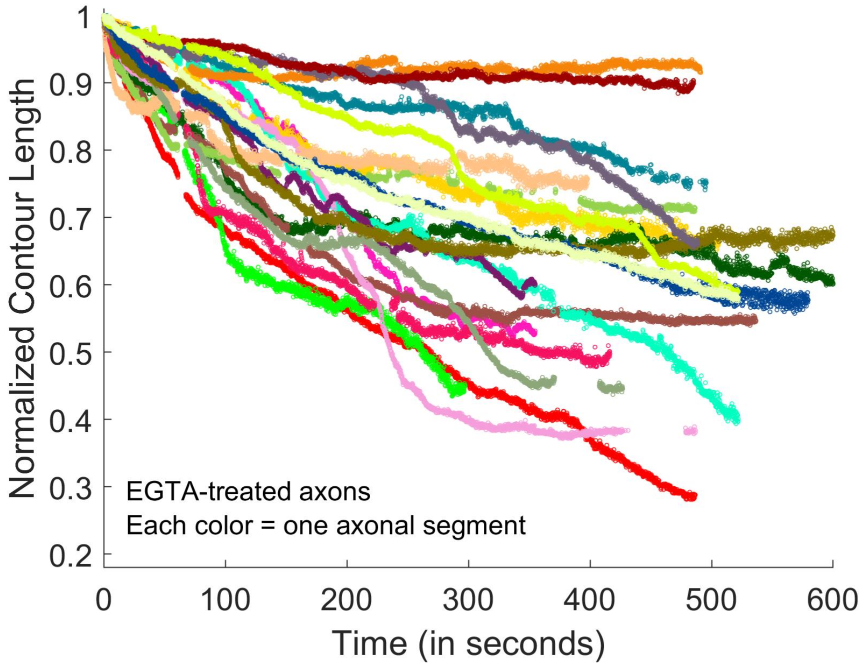


**Fig. S5:** Normalized contour length is plotted against time for individual axonal segments maintained in medium containing 5 mM EGTA. Each color represents one segment. These axons retracted only partially and did not recover within the observation period of approximately 600 s. The gap near t $\approx$ 57 s results from switching the recording rate from 15 to 6 fps.

#### Microtubule and axonal caliber recovery in $Ca^{++}$-deficient condition.

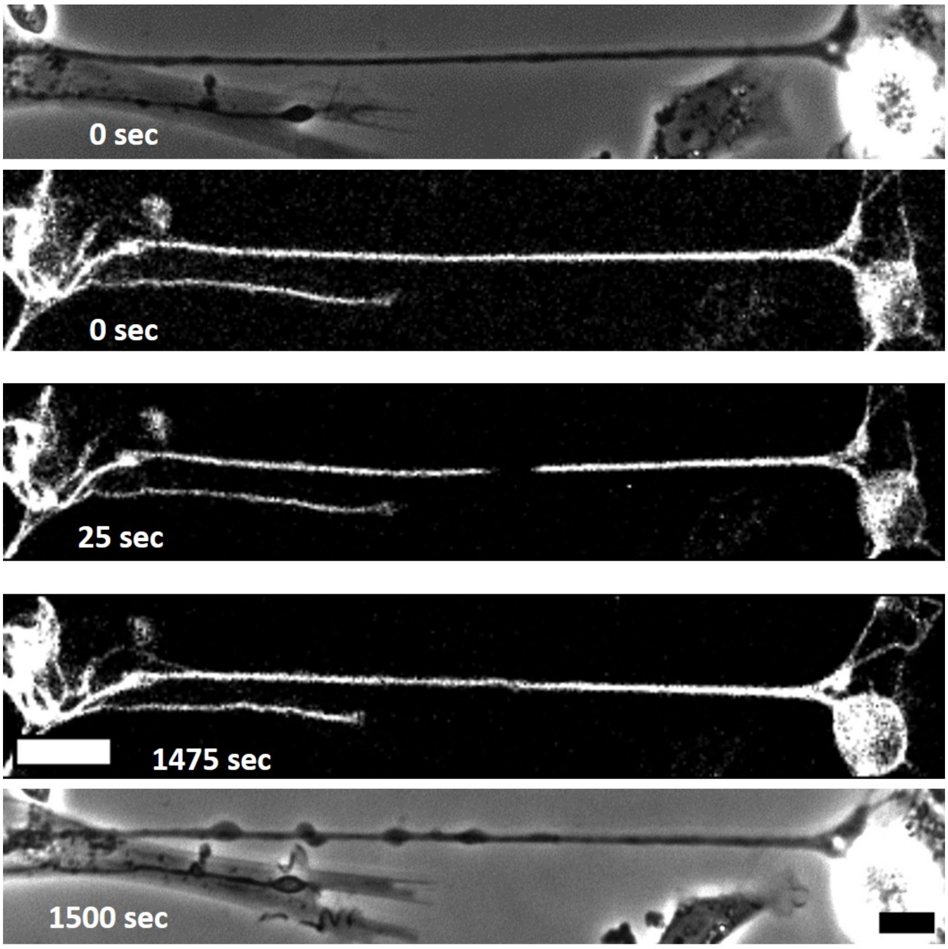


**Fig. S6:** Phase contrast and fluorescence image sequence of an axon labelled with a microtubule-specific dye (SPY555-Tubulin) under calcium-deficient conditions. Images at 0 s represent the pre-ablation state in both phase contrast and fluorescence. The fluorescence image taken at 25 s post-ablation shows the partial retraction of microtubules in the same axon. Images taken at 1475 and 1500 s show the extent of recovery in microtubule intensity and axonal caliber, respectively, within the damaged region. The scale bars for the fluorescence and phase contrast images are 20 $\mu$m and 10 $\mu$m, respectively.

**Kymograph of a control axon transfected with the pCAG-mNeon-EB3 construct.**

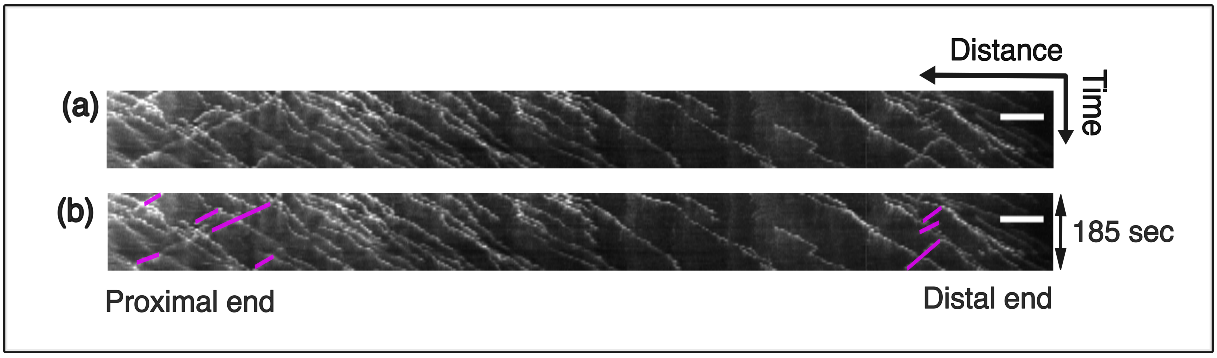


**Fig. S7:** (a) Kymograph showing EB3 comets moving predominantly in the anterograde direction, indicating a uniformly plus-end-out microtubule orientation. Only a few retrograde comets are present, indicated by the magenta colour in (b). Scale bar: 5 $\mu$m.

**Kymograph of axons transfected with the pCAG-mNeon-EB3 showing resealing event in $Ca^{++}$ chelated media.**

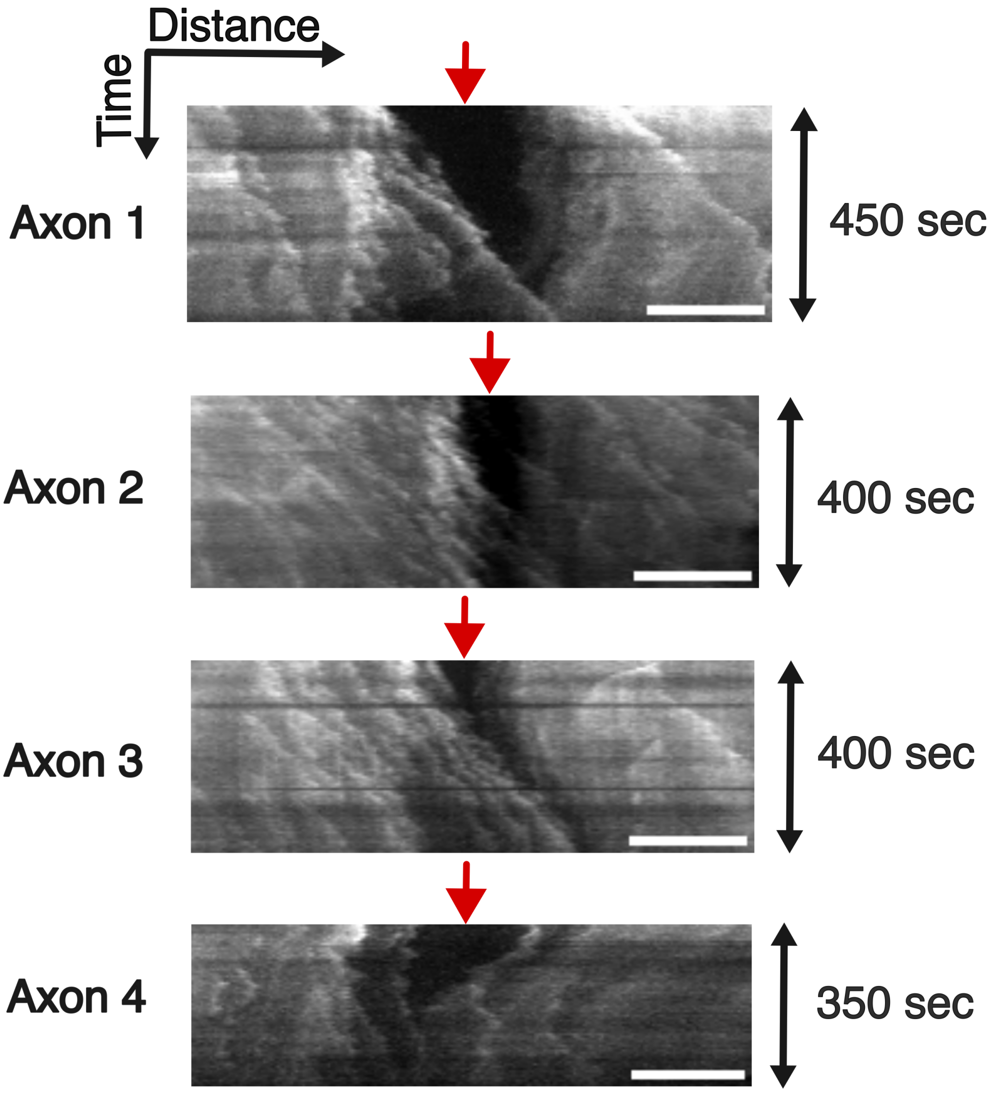


**Fig. S8:** Kymographs of partial or complete resealing events for four different axons that were maintained in $Ca^{++}$ chelated media. The injury sites are marked by red arrows. Higher comet invasion occurs from the cell body side(left) in the first three cases. Scale bar: 5 $\mu$m.

Supplementary Table S1

**Statistical analyses for Figures 3c and 8.** Normality was assessed using the Shapiro–Wilk test. Non-parametric tests were used because at least one group in each analyzed dataset did not satisfy the normality assumption.

| Figure | Groups and sample size | Normality test | Statistical test | Result |
|---|---|---|---|---|
| Figure 3c | Control, $n = 40$ axons; EGTA, $n = 47$ axons; BAPTA-AM, $n = 9$ axons | Shapiro–Wilk: Control, $W = 0.83$, $p < 0.0001$; EGTA, $W = 0.99$, $p = 0.993$; BAPTA-AM, $W = 0.93$, $p = 0.508$ | Kruskal–Wallis test followed by Dunn's multiple-comparison test | Kruskal–Wallis: $H(2) = 76.69$, $p < 0.0001$. Dunn's test: control vs EGTA, $p < 0.0001$; control vs BAPTA-AM, $p = 0.047$; EGTA vs BAPTA-AM, $p = 0.019$. |
| Figure 8 | F-actin, $n = 32$ axons; microtubule, $n = 11$ axons | Shapiro–Wilk: F-actin, $W = 0.86$, $p < 0.001$; microtubule, $W = 0.95$, $p = 0.678$ | Two-tailed Mann–Whitney U test | $U = 350$, $p < 0.0001$. |

#### Captions for video files from Video 001 to 007

**Video 001:** Time-lapse recording of an axon after partial ablation. The video captures the pre-ablation state, the ablation event, and subsequent retraction dynamics toward the proximal and distal ends. Over time, the cytoskeleton, along with the cytoplasm, retracts toward the extremities, keeping the membrane intact.

**Video 002:** Time-lapse recording of an axon following full ablation, showing the axon splitting into two segments after laser injury. The recording captures the pre-ablation frame, the ablation event, and subsequent retraction evolution toward both ends. Note that the axon buckles heavily and the two cut ends are not aligned as in the case of partial ablation.

**Video 003:** Time-lapse recording of a paraformaldehyde-fixed axon following full ablation. As shown in the video, the axonal segments do not retract along the axonal axis after ablation. This indicates that the retraction dynamics observed in Video 001 are intrinsic to the axon and not a result of the laser impact.

**Video 004:** Time-lapse recording of an axon treated with a $Ca^{++}$ dye, Fluo-4 AM, showing calcium elevation following partial ablation.

**Video 005:** Time-lapse recording of an EGTA-treated axon stained with a microtubule dye, showing microtubule recovery after partial ablation.

**Video 006:** Time-lapse recording of an axon transfected with pCAG-mNeon-EB3 construct. The video has been gray-scale inverted using Fiji LUT to improve clarity. It can be seen that microtubule regrowth occurs after partial ablation and an initial retraction when axons are maintained in $Ca^{++}$ free media. The cell body is on the left.

**Video 007:** Time-lapse recording of an EGTA-treated axon stained with a filamentous actin dye, showing actin filament recovery after partial ablation.
